# Varing chemical equilibrium gives kinetic parameters

**DOI:** 10.1101/000547

**Authors:** Edward Flach, Santiago Schnell

## Abstract

We are interested in finding the kinetic parameters of a chemical reaction. Previous methods for finding these parameters rely on the dynamic behaviour of the system. This means that the methods are time-sensitive and often rely on non-linear curve fitting.

---

In the same manner as previous techniques, we consider the concentrations of chemicals in a reaction. However, we investigate the static behaviour of the reaction at dynamic equilibrium, or steady state. Here too, the chemical concentrations depend on the kinetic parameters of the reaction. In an open reaction, the static concentrations also depends on the rate of input of the source of reacting chemical. Controlling this input rate slides the steady state along a curve in concentration space. This curve is determined by the kinetic parameters. The plane of this curve is sufficient to find the kinetic parameters.

The new method we propose uses only the steady state concentration values to determine the kinetic parameters of the reaction. These values are constant once dynamic equilibrium is achieved, and so can be read accurately. Readings can be repeated readily to reduce error. Thus this new technique is simple and could produce accurate kinetic parameter estimates.

## 1 Finding rate parameters in closed systems

Chemical reactions have shape, which is formalised as stoichiometry, and speed, which is given by rate parameters. The first step is to find the stoichiometry; here we are interested in the second step. There are four general approaches for finding these rate parameters [2, 4]:

1. *transient kinetics* reaction behaviour is tracked during the initial fast transient
2. *initial rate* reaction behaviour is tracked after the initial fast transient
3. *progress curve* reaction behaviour is tracked during the secondary slow transient
4. *relaxation* reaction behaviour is tracked as it approaches equilibrium

These methods are based on an understanding of the reaction progress curve. In each case, assumptions are made such as, for the initial rate method, that the substrate concentration remains at initial levels. These simplifying assumptions allow analysis, and perhaps require measurement of fewer species.

The reaction progress curve has been broken down and characterised according to characteristic behaviours in each stage of the reaction. This characterisation is reasonable, however a basic assessment of reaction time scales must be made before these techniques can be attempted. In each case, the assessment of time scales combined with the assumptions gives the possibility of errors.

We propose a new method for determining reaction rates. Our approach gives the natural completion to the breakdown of the reaction curve: we stop at the end. We are only interested in the steady state, the final destination of the reaction curve. The is also known as chemical equilibrium.

Our technique works by determining the steady state in multiple circumstances. We require a significant change in circumstances, and for that the reaction must be open. We can then change the steady state by altering the input to the reaction. As the steady state varies, the kinetic parameters are revealed.

This is not a standard perturbation. In fact, the term “perturbation” is not strictly correct: we do not perturb the system from its steady state, we perturb the steady state itself. We vary a component of the system, and the steady state moves in response. However, this movement is controlled; the steady state is determined by the rate constants of the system.

This approach is novel and practical. Since only the steady state data is required, the method is robust: standard methods require dynamic data, which is more sensitive to measurement.

## 2 Open reactions are very different to closed

A closed reaction is one where all the reactants are present at the start. There is no external variation to the system as the reaction occurs. The reaction runs to completion. This is the normal type of reaction of in vitro experiment.

In contrast, an open reaction is the type that occurs in vivo. Here there is continuous supply, or source, of some reactant. There is also removal, or sink, of some product. The supply need not be constant, and in general would not be. However, here we consider only the simplest form: a constant supply. The strength of the source is important; it has a substantial effect on the reaction dynamics.

### 2.1 What does it mean to have a constant supply?

Mathematically, this is simple. If the reactant or substrate is denoted S, then is has concentration [*S*], but we write this more simply as *s*. If there is no reaction then we can say that the change in concentration d*s/*d*t* = *µ*, constant.

### 2.2 What does constant supply mean experimentally?

We wish to keep the reacting volume constant, since otherwise we effectively change the concentration of the other chemicals involved in the reaction, altering the reaction rates. Therefore, the obvious situation is to supply substrate in solid form, at a constant rate. This is not always possible. Furthermore, solid substrate might dissolve unevenly, effectively causing fluctuations in the supply.

Therefore we consider a steady physical trickle of substrate as a high-concentration solution. We assume that the change in volume will be negligible. An alternative method of supply could be via membrane diffusion. In this case, we have a reservoir with a high concentration of source reactant. The membrane then supplies this reactant at a fixed rate through the membrane. We require no other transfer through the membrane.

A third method of substrate supply would be a pre-substrate chemical which supplies the substrate. If this reaction is irreversible, and the pre-substrate clamped at a high level, the substrate could be supplied at a constant rate. Since the pre-substrate chemical would be present in the mix at the start, then the solvent volume will remain fixed.

At the end of the reaction, we require removal of the product. This is known as a sink. This is given if the product is insoluble or a gas. Otherwise we need to ensure that the product is removed. An irreversible reaction is required, and the co-reactant should be heavily in excess (clamped). In this case a constant steady state can be reached.

We consider how this system differs from a closed system. If the sink is removed, then the system will “blow up” since there is a constant net supply to the system. That is, the concentration of one or more of the reactants will increase continuously. If the source is removed, the system will drain, leaving it at the zero steady state. Thus both a source and a sink are required for a viable open system. When both source and sink are removed we have a closed system.

## 3 A simple example reaction

We consider a single-enzyme, single-substrate reaction. The elementary step here is substrate *S* binding to enzyme *E* to form a complex *C*. The enzyme may effect a change in the substrate, and after some time will dissociate. Thus the enzyme is returned unchanged, and the substrate has turned into the new product molecule *P*. Each step in this process could potentially be reversible. The source and sink must however be irreversible. We show this diagrammatically:

Forward formation of complex has parameter *a*, and disassociation *b*. The corresponding *α* and *β* indicate the rates of the reverse reaction steps. Here we have indicated the constant source term *µ* and a sink rate *m*.

Using the law of mass action, we convert this scheme into a set of differential equations:

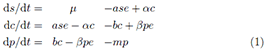

We choose example rate parameters and simulate the reaction course using COPASI, which is a tool for simulation and analysis of biochemical pathways. Its predecessor, GEPASI [6, 7, 8], is widely used to test new parameter estimation techniques on computers. We select the stochastic method for a very small test volume, and run until the system appears steady. The time-course of the concentrations is shown for one of these simulations in figure 2.

**Figure 1:**
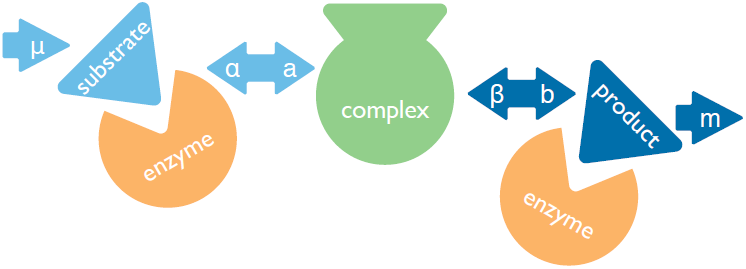
A simple example reaction: enzyme catalysis. The Michaelis-Menten reaction starts with a substrate binding to an enzyme. The enzyme assists the substrate to convert to the product, Each step is reversible, the first step has forward rate *a* and reverse rate *α*. Th second step has forward rate *b* and reverse rate *β*. This is an open system with a supply of substrate at rate *µ* and removal of product at rate *m*.

**Figure 2:**
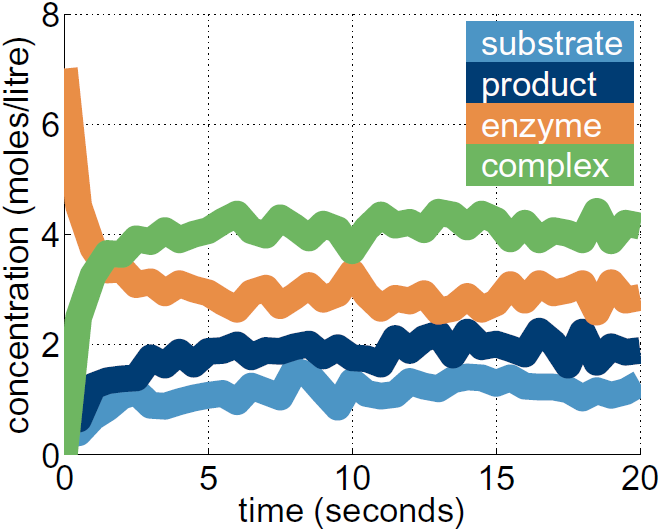
Simulation of example reaction shown in figure 1 using COPASI. The experiment is run until the system steadies. Here the source flow *µ* is 4 mole/litre/second, *a* = 6, *b* = 5, *α* = 4, *β* = 3, *m* = 2. Initial (total) enzyme concentration is 7 moles/litre, and product and substrate is 1.

Initially there is rapid change in the concentrations of the reactants. After some time the reactions settle down: this means we have reached a steady state. Once this has happened, we can determine the values for this steady state. Given the reaction in figure 2, we take readings from 10 seconds onwards. This gives an estimate of the steady state values.

## 4 Reactions progress to chemical equilibrium

There is a concept of chemical equilibrium [9] which occurs when the flux (the net flow of the reaction) of a system is zero. In a closed system, this is equivalent to the steady state. A system at chemical equilibrium is at steady state. However, a system at steady state does not have to have zero net flow, and so may not be at chemical equilibrium. In an open system, we have a net flux through the system, even at the steady state. A zero flux occurs when the source is removed, producing a zero steady state. So the concept of chemical equilibrium is not useful in open systems. To make the distinction one might refer to non-equilibrium steady states [10]. This essential difference between open and closed systems is why standard techniques used in closed systems do not always apply to open systems.

We consider the fixed point of the system, or steady state. In this situation all the derivatives are set to zero: d*s/*d*t* = d*c/*d*t* = d*p/*d*t* = 0. This reduces the problem to an algebraic one:

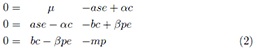

We manipulate the equations. We see that this produces a cascading effect where the flow is equalised through the system:

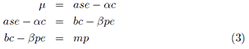

We can see this effect more clearly when we write these equivalences together

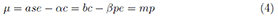

If we compare this to the reaction, figure 1, we see that each term is the flux through each stage of the reaction: the arrows *between* the chemical species, or states of the reaction.

We consider the start and end flux terms, namely *µ* = *mp*. This tells us that, at steady state, the flow into the system equals the flow out. In general we see that, at steady state, the flow is constant and equal between each state.

## 5 Varying equilibrium gives rate parameters

We find the concentrations of all the reactants at the steady state. Then we can use the set of equivalences we found in equation 3. We consider the components of the equations, the reaction velocities such as *ase* or *αc*. Now we introduce some new terminology: the rate function. We define this as the variable part of the reaction velocity (such as *se* or *c*), with the rate constant (such as *a* or *α*) the constant part. The product of the rate constant and the rate function is the rate of the reaction.

We consider the first steady state equation, *µ* = *ase − αc*. Here *a* is the rate constant, with the corresponding rate function *f* (*s, e*) = *se*. The reverse reaction has a simpler rate function: *g*(*c*) = *c*. The next step is to consider the rate functions as variables: *x* = *f* (*s, e*) and *y* = *g*(*c*). The final step is to consider the source rate as a variable too; we define *z* = *µ*. Then the equivalence becomes *z* = *ax − αy*. This is the equation of a plane. The parameters of the plane are *a* and *α*, the rate constants we are interested in.

We alter the source rate *µ*. For each value of *µ* we find corresponding values for each chemical species at steady state. From this, we calculate the rate function variables *x*, *y* and *z*. We find the plane that these data points lie on, shown in figure 3. For data fitting we use Singular Value Decomposition. An alternative method is Principal Components Analysis. These are least squares fit to the data. We plot the plane on which the (*x, y, z*) data points lie, and from this find values for *a* and *α*. The formula found for the plane is *z* = 6.04*x −* 4.02*y*. This is close to the original values of the simulation, namely *a* = 6, *α* = 4.

**Figure 3:**
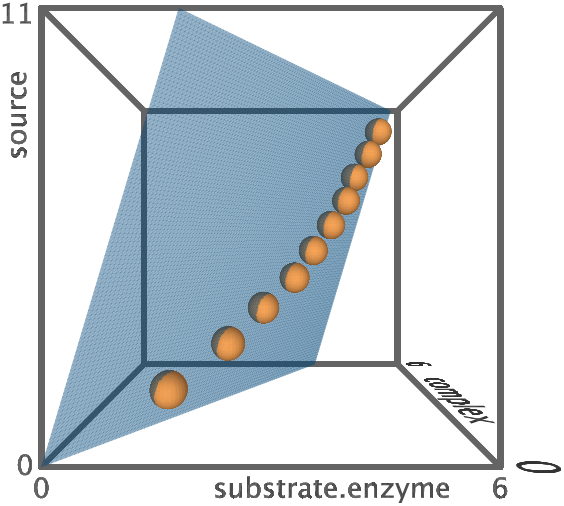
The steady state slides along a curve. This curve sits in a plane which we use to determine reaction parameters. Data is for a variable source term, *µ* = 1–10, in unit steps. The plane is found: *z* = 6.12*x* 4.12*y*, which corresponds to the reaction rates of the simulation: *a* = 6, *α* = 4 in *z* = *ax* − *αy*.

We can repeat this process for the second steady state equation, yielding estimates for *b* and *β*. The third steady state equation, *µ* = *mp*, gives a straight line. Fitting the data in this case is far simpler.

The plane given always runs through the origin, so we know one point before we begin. This precise data point improves the accuracy of the technique.

This result comes from observing that the steady state value for each reactant is dependent on the source and sink of the system. This dependency is non-linear. The reduction of the non-linear terms to simple variables gives a linear relationship between these rate functions and the source rate.

## 6 How much to vary; how many replicates?

For this method, we must repeat the experiment for several source levels. In our example, *µ* = { 1, 2 … 10}. For each fixed source level we could repeat the experiment several times.

We consider the accuracy of the parameter determination. Since we chose the parameters for our numerical experiment, we can compare our results to the actual values. We calculate the percentage error of each parameter found in an experimental configuration, and consider the worst of these – the maximum error (as shown in figure 4). We see that increasing the number of steady state points (distinct input flux levels) improves results more efficiently than repeating with the same flux levels.

**Figure 4:**
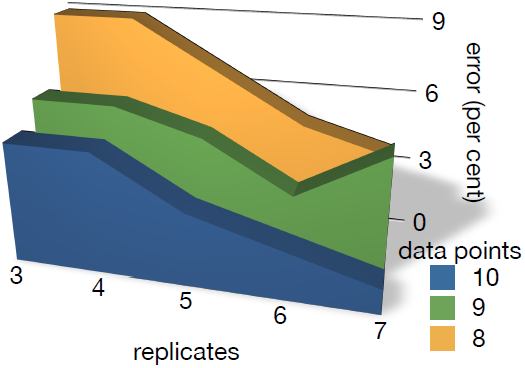
Accuracy of experimental approaches. The graph shows the worst error overall as a percentage of the original parameter value (for *a*, *α*, *b*, *β* and *m*). Each different input flux (*µ* = constant) run to steady state gives one point. We see that increasing the number of points improves accuracy more than repeating the experiment with the same flow.

There is a good reason why this should be the case. We can see clearly in figure 2 that the points lie on a curve. This curve lies in a unique plane, but to find the plane accurately the curve has to be distinctly, well, curved. In contrast, if the points lie roughly on a straight line then many different planes can fit the data and our parameter estimation will be poor. The more flux levels we can sample from, and the bigger their range, the more likely there will be a good distribution. Then the plane can be found accurately. The formula for the plane directly gives the kinetic parameters and so these will then be good estimates.

## 7 The new method is useful

We have shown that the system behaviour at steady state can be used to determine every rate constant in a simple reaction. This approach does rely on prior knowledge of the reaction mechanism and kinetics. It also relies on the ability to read all relevant concentration levels, but only their value at steady state is required.

By simulating an experiment, we have seen that the approach is viable. With a limited number of data points and few repeats we produce a high level of accuracy. The most significant observation for experimental design is that the broader the range of input levels, the higher the accuracy of the data fit. Maximising this variation in flux levels is the best guarantee of good results.

Previous theory inspired by the availability of high-throughput technology has emphasised the need to measure the time-courses of chemical concentrations [3], or the reaction velocities. This new technique is far more simple – seemingly naively so. By only requiring the static values of the chemical concentrations at their steady states we suggest an opposite requirement to previous analyses. Hopefully, this data will be far easier to produce experimentally, since the time-dependence and transient behaviour is not required.

Our basic premise is that the system will eventually reach a steady state. This does not always happen [11]. However, in the general case, the steady state is often stable [12].

The method is straightforward and experimentally robust. This is a strong result gained through a simple technique. Now it is for an experimentalist to find a method for carrying out the experiment practically.

## Acknowledgements

We would like to acknowledge support from NIH grant number R01GM076692.

